# Population-level consequences of inheritable somatic mutations and the evolution of mutation rates in plants

**DOI:** 10.1101/2020.09.29.318402

**Authors:** Thomas Lesaffre

## Abstract

Inbreeding depression, that is the decrease in fitness of inbred relative to outbred individuals, was shown to increase strongly as life expectancy increases in plants. Because plants are thought to not have a separated germline, it was proposed that this pattern could be generated by somatic mutations accumulating during growth, since larger and more long-lived species have more opportunities for mutations to accumulate. A key determinant of the role of somatic mutations is the rate at which they occur, which likely differs between species because mutation rates may evolve differently in species with constrasting life-histories. In this paper, we study the evolution of the mutation rates in plants, and consider the population-level consequences of inheritable somatic mutations given this evolution. We show that despite substantially lower per year mutation rates, more long-lived species still tend to accumulate larger amounts of deleterious mutations because of higher per generation, leading to higher levels of inbreeding depression in these species. However, the magnitude of this increase depends strongly on how mutagenic meiosis is relative to growth.

## 1 Introduction

Plant growth is fueled by cell divisions occuring in meristems. Each shoot is produced by an apical meristem and may bear axillary meristems, which are typically situated in the axils of leaves and grow out to become the apical meristem of a new shoot upon activation (Burian et al., 2016). As meristematic cells generate all the tissues constituting the shoot, any mutation occuring in a meristematic cell will be borne by all the cells it gave rise to, leading to genetic mosaicism within individual plants. Furthermore, because meristems also give rise to reproductive tissues, mutations occuring during growth before the differentiation of the germline, that is somatic mutations, may be present in the gametes and hence be inherited (Lanfear, 2018). All else being equal, it follows that the larger and the older a given plant grows, the more somatic mutations it should accumulate and transmit to its offspring, potentially leading to a higher mutation load in more long-lived and larger species since it is thought that most mutations are deleterious (Eyre-Walker and Keightley, 2007).

Inbreeding depression, that is the decrease in fitness of inbred relative to outbred individuals (Charlesworth and Charlesworth, 1987), is thought to be mostly generated by recessive deleterious mutations maintained at mutation-selection balance in populations (Charlesworth and Willis, 2009). Hence, Scofield and Schultz (2006) proposed that somatic mutations accumulation could lead to higher inbreeding depression in larger and more long-lived species. Consistent with this view, inbreeding depression was indeed shown to increase strongly as life expectancy increases across plant species (Duminil et al., 2009; Angeloni et al., 2011). Furthermore, Bobiwash et al. (2013) showed that substantial in-breeding depression was generated by somatic mutations in a study performed at the phenotypic level in old *Vaccinium angustifolium* clones. This is however, to our knowledge, the only empirical test of Scofield and Schultz (2006)’s. Besides, recent theoretical investigations have shown that variations in inbreeding depression can in principle be generated by differences in the fitness effect of mutations between species with contrastring life-histories (Lesaffre and Billiard, 2020), so that somatic mutations accumulation may not always be needed to explain variations in the magnitude of inbreeding depression across plant species. Moreover, theoretical investigations of the population-level consequences of somatic mutations accumulation are lacking, so that their role in the maintenance of high inbreeding depression in long-lived species remains poorly understood. Indeed, theoretical studies regarding somatic mutations in plants either focused on the case of favorable mutations, conferring resistance against herbivores (e.g. Antolin and Strobeck, 1985), or studied the fate of deleterious mutations subject to intra-organismal selection (Otto and Orive, 1995; Pineda-Krch and Lehtilä, 2002), but never considered the population-level consequences of recessive deleterious mutations (Schoen and Schultz, 2019). In summary, deleterious somatic mutations accumulation has been proposed as a mechanism to explain the rarity of selfing species among long-lived plants (Scofield and Schultz, 2006), consistent with empirical measures of inbreeding depression, but theoretical support for this idea remains scarce.

An important determinant of the consequences of somatic mutations accumulation is the rate at which said mutations accumulate during growth, that is the somatic mutation rate, which is defined here as the number of mutations occuring per unit of vegetative growth. This rate is likely influenced by evolutionary mechanisms similar to those affecting mutation rates in general. For example, Kimura (1967) showed that mutation rates should be shaped by the opposition between the increase in the number of deleterious mutations borne by individuals with higher mutation rates on the one hand, which causes indirect selection against genetic variants increasing mutation rates to increase, and the direct fitness cost there is to increasing the fidelity of DNA replication on the other hand. Besides, Lynch (2011) proposed that selection to decrease the mutation rate should become weaker than genetic drift at some point in finite populations, thereby favoring the persistence of non-zero mutation rates. Nevertheless, the inheritability of somatic mutations in plants and their intrinsic link with growth and life expectancy likely contribute to shape the evolution of mutation rates in a specific manner which was, to our knowledge, never tackled theoretically. Great interest was however taken in empirically detecting somatic mutations and comparing mutations rates in a variety of plants species ranging from the very short-lived *Arabidopsis thaliana* to ancient, centuries old trees. In an analysis performed across many plant families, Lanfear et al. (2013) showed that taller species among pairs of sister species have signficantly lower rates of molecular evolution, measured as the number of substitutions per site per 10^6^ years. They argued that contrary to animals, this pattern is not a mere reflection of differences in generation time, which would reflect different rates of genome copying per unit of time, because somatic genome copying events contribute to the inheritable genetic variation in plants. Instead, they proposed that this pattern may be due to slower growth in taller species, which results in a lower number of mitosis (and therefore mutations) per unit of time. Consistent with this view, it was shown at the cellular level that axillary meristems cells are set aside early during the growth of a shoot (Burian et al., 2016), resulting in a number of cell divisions increasing linearly with the number of branching events in trees although the number of terminal branches increases exponentially. Furthermore, multiple studies showed that somatic mutation rates tend to be considerably lower in taller, more long-lived species (Schmid-Siegert et al., 2017; Plomion et al., 2018; Hofmeister et al., 2019; Orr et al., 2020; Wang et al., 2019; Hanlon et al., 2019). For instance, Orr et al. (2020) found the somatic mutation rate per generation to be only ten times higher in *Eucalyptus melliodora* than in *Arabidopsis*, despite being *>* 100 times larger in size.

Thus, empirical evidence indicates that more long-lived species have acquired mechanisms to reduce the amount of mutations accumulated during growth on the one hand, but still present high levels of inbreeding depression on the other hand, which suggests that more long-lived species still accumulate more mutations despite above mentioned limiting mechanisms. The aim of the present study is to disentangle the relationship between these two observations. We first study the evolution of the mutation rate in plants, and then consider the number of mutations and the magnitude of inbreeding depression maintained at mutation-selection balance, given the evolutionarily stable mutation rate reached by the population. To do so, we extend the work of previous authors (Kimura, 1967; Gervais and Roze, 2017) to the case of a perennial population in which individuals grow as they age and accumulate mutations in doing so. We obtain analytical predictions which we test against the output of individual-centered simulations. We show that the evolutionarily stable mutation rate should decrease in plants as life expectancy increases, because deleterious mutations have more time to accumulate in more long-lived species. Furthermore, we show that despite substantially lower per year mutation rates, more long-lived species still tend to accumulate larger amounts of deleterious mutations because of higher per generation, leading to higher levels of inbreeding depression in these species. However, the magnitude of this increase depends strongly on how mutagenic meiosis is relative to growth.

## 2 Methods

### Model outline

We consider a large population of hermaphroditic diploids. Individuals survive between mating events with a constant probability *S*. Juveniles may only settle in replacement of deceased individuals, so that population size is kept constant. We assume that individuals are made of a trunk, which grows by one section between each flowering event.

In our model, mutations at the selected loci occur both during meosis and somatic growth. The somatic mutation rate per unit of growth, that is per new section produced (*u*), of a given individual is determined by its genotype at a single modifier locus. At this locus, we consider the fate of a rare mutant (*m*) with a weak effect (*ε*) competing with a resident allele (*M*). We assume this mutant allele to be codominant with the resident, so that an individual’s somatic mutation rate is given by *u*_*MM*_ = *u*_0_, *u*_*Mm*_ = *u*_0_ + *ε*, or *u*_*mm*_ = *u*_0_ + 2*ε*, depending on its genotype at the modifier.

Although meiotic and somatic cell divisions likely share common features, so that meiotic and somatic mutation rates should evolve together to some extent, they also differ in various ways. For instance, recombination during meiosis causes additional breaks in DNA strands, which gives the opportunity for more mutations. Furthermore, somatic and meiotic mutation rates are not defined on the same scale. Indeed, while the somatic mutation rate is usually defined as a number of mutations per unit of growth, meiotic mutation rates are defined at the scale of a whole reproductive event. Thus, the relationship between these two mutation rates is not straightforward, because different genetic events happen and divisions occur at different paces. For the sake of simplicity, we will assume that meiotic mutations are produced at rate *γu*, where *γ* is a positive real number which allows one to tune the intensity of meiotic mutation relative to somatic mutation. In other words, we assume there is a linear relationship between the two rates.

We assume that any section can contribute to reproduction (Fig. 1). Self-fertilisation occurs at rate *α*, a fraction *σ* of which imperatively occurs within the same section. The remaining fraction 1 − *σ* can occur between sections within the individual. A section’s fecundity is determined by its genotype at a very large number of biallelic loci acting multiplicatively. At these loci, allele 0 is an healthy allele, while allele 1 is a mutated allele which diminishes the section’s fecundity by a proportion *s*. In heterozygotes, allele 1 expresses proportionally to its dominance coefficient *h*. Following previous authors (Gervais and Roze, 2017), we also introduce a DNA replication fidelity cost function, *f*, which is an increasing function of the meiotic mutation rate *γu*. Gervais and Roze (2017) considered a variety of cost functions and came to qualitatively similar conclusions in every case. Yet, most of their results were obtained using the cost function given in Equation (1),

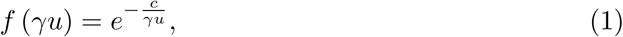

where *c* is the cost of replication fidelity, which we also use in this study. Thus, the fecundity of a section is given by

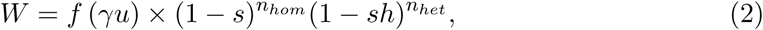

where *n*_*hom*_ and *n*_*het*_ are the number of mutations borne in the homozygous and heterozygous states, respectively.

**Figure 1:**
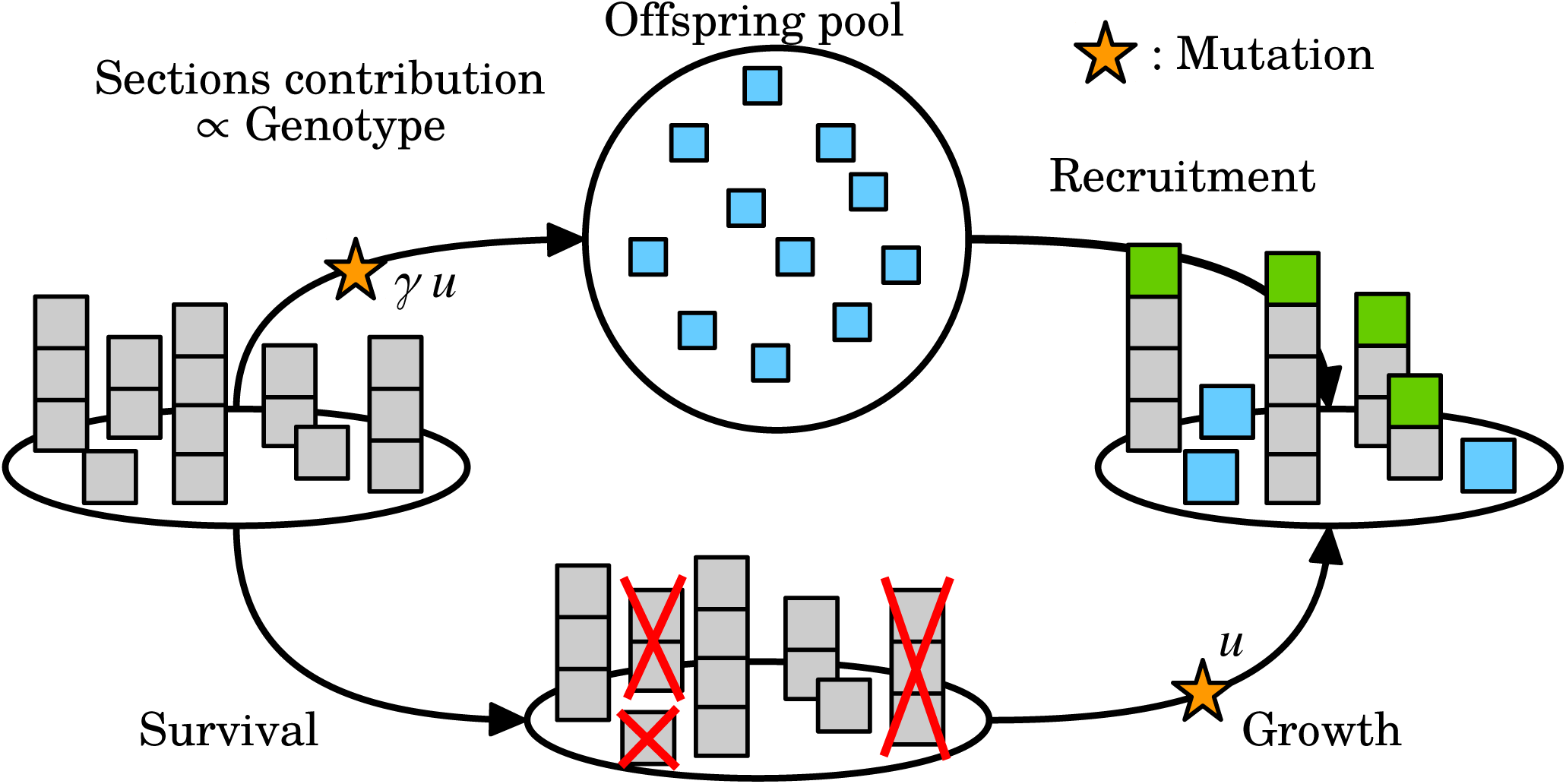
Life cycle of the modeled population. Blue squares are offspring, which are made of a single section. Green squares depict the sections gained during growth by survivors. Yellow stars indicate the steps at which mutation occurs.

### Analytical methods

We use the theoretical framework described in Kirkpatrick et al. (2002) to study our model, which relies on indicator variables to describe individuals’ multilocus genotypes. In the analytical model, we neglect the effect of the proportion of obligate within-section selfing (*σ*) since it will prove to have very little impact on our results. For the sake of brevity, derivations of the results presented in the following sections are detailed in Appendices I.1 and I.2 for results regarding the evolution of mutation rate and the mutation-selection equilibrium properties of the population given the evolution-arily stable mutation rate, respectively.

### Individual-centered simulations

We ran individual-centered simulations to test the validity of our analytical approximations. The simulation program was coded in C++11 and is available from GitHub (https://github.com/Thomas-Lesaffre/Somatic_mutations). In this program, individuals are represented by two chromosomes of length *λ* (expressed in cM) with the modifier situated at the center and along which mutations can occur at any position, so that infinitely many selected loci are effectively modeled (Roze and Michod, 2010).

### Modeled loci

At the modifier, we assume that infinitely many alleles exist, coding for any value of *u* ∈ [0, +∞[. Mutation occurs at rate *u*_*m*_ = 10^*−*3^, and the value coded by the new allele is sampled from a Gaussian distribution centered on the former allele value with standard deviation *σ*_*m*_ = 10^*−*2^, which is truncated at zero to prevent the modifier from going out of range. At selected loci, the number of mutations occuring on a chromosome during a given mutation event is sampled from a Poisson distribution with mean *u* (*γu* for meiosis), and their position is sampled from a uniform distribution. Recombination is modeled by exchanging segments between homologous chromosomes. The number of crossing-overs is sampled in a Poisson distribution with mean *λ* and their positions are sampled from a uniform distribution along chromosomes. Every time a mutation occurs, the age of the section at which it occured along the individual is stored, so that the genotype of any section within an individual can be reconstructed at any time from the individual genome. This method allows us to gain substantial computation time because mutations are stored only once per individual instead of being copied once for each new section.

### Sequence of events

The population is kept of constant size, *N*. Between each mating event, individuals have a constant survival probability *S*. If they survive, they grow by one section, and mutations occur at rate *u* in this section. If they die, they are replaced by an offspring produced by the population. Any section within any individual can be chosen as a parent, with a probability proportional to its fecundity (Equation 2). The offspring is produced by self-fertilisation with probability *α*, in which case the chosen section mates with itself with probability *σ*, and with any section within the same individual with probability 1 − *σ*. When selfing occurs between sections, a second parental section is selected within the individual. When the offspring is not produced by self-fertilisation, which occurs at rate 1 − *α*, it is produced by random mating and a second parent is selected from the whole population. Mutation occurs at rate *γu* during meiosis.

### Measurements

Once the equilibrium was reached, that is when both the mutation rate and the average number of mutations per chromosome were at equilibrium, we measured the average number of mutations per chromosome in seeds, the average mutation rate and inbreeding depression. Although individuals are chimeric in our model, we stuck to measuring inbreeding depression at the individual level to be in line with its definition. To do so, we counted how many times each individual was chosen as a parent before it died (*i*.*e*. we measured its lifetime reproductive success) and used this quantity as a measure of lifetime fitness. Individuals were marked as being produced by outcrossing (0), selfing within the same section (1), and selfing between sections within the same individual (2), so that we were able to measure fitness differences between these various categories of individuals. Namely, we measured inbreeding depression, that is the decrease in fitness of selfed individuals relative to the outcrossed (*δ*_01_), and autogamy depression (Schultz and Scofield, 2009; Bobiwash et al., 2013), that is the decrease in fitness of within-section selfed individuals relative to between-sections ones (*δ*_12_). Ten replicates were run for each parameter set. Simulations were kept running for 10^6^ and 2 × 10^5^ reproductive seasons for life expectancies lower and higher than 200 reproductive seasons, respectively. Results were averaged over the last 10^5^ reproductive cycles (resp. 2 × 10^4^) and the 95% confidence interval around the mean was also recorded.

## 3 Results

In what follows, life expectancy (*E*) will be used to discuss results instead of survival probability (*S*) for the sake of clarity and biological relevance. Given survival probability *S*, life expectancy can be computed as

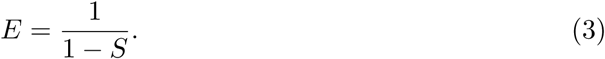

### 3.1 Evolutionarily stable mutation rate

Let us first study the evolution of the mutation rate. We show in Appendix I.1 that the evolution of the mutation rate is the result of the opposition between the direct cost of DNA replication fidelity, which is higher when the mutation rate is lower, and the indirect selection caused by deleterious alleles which tend to be more frequently linked with modifier alleles increasing the mutation rate (Equation A23). The resulting evolutionarily stable mutation rate is given by

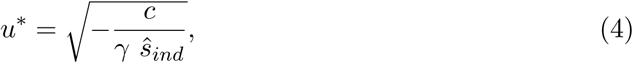

where *ŝ*_*ind*_ encapsulates the intensity of indirect selection acting on the modifier. Its expression is derived in Appendix I.1.5. Figure 2 shows the evolutionarily stable mutation rate as a function of life expectancy (top row), along with the intensity of indirect selection (bottom row), for cases where *γ* = 1, *γ* = 10 and *γ* = 100. We chose to focus on cases where *γ* ⩾ 1, that is on cases where more mutations are produced during meiosis than during the development of a new section, on the basis of three lines of evidence. First, direct observations of plant development at the cellular level indicate that cells destined to form axillary meristems undergo much fewer divisions than other cells from the moment they are produced in the apical meristem, which suggests that the number of cell divisions per branching event, and therefore the number of opportunities for mutations to accumulate, may be lower than previously thought (Burian et al., 2016). Second, estimates of somatic mutation rates per unit of growth tend to be low (Orr et al., 2020). Third, to our knowledge, the only experiment comparing the mutagenicity of meiosis and mitosis was performed by Magni and Von Borstel (1962) in yeast. They found meiosis to be 6 to 20 times more mutagenic than mitosis, which further suggests that *γ* may tend to be greater than 1. Besides, performing simulations *γ <* 1 proved to be very challenging since the number of mutations accumulated in the population quickly became very high, causing simulations run very slowly.

**Figure 2:**
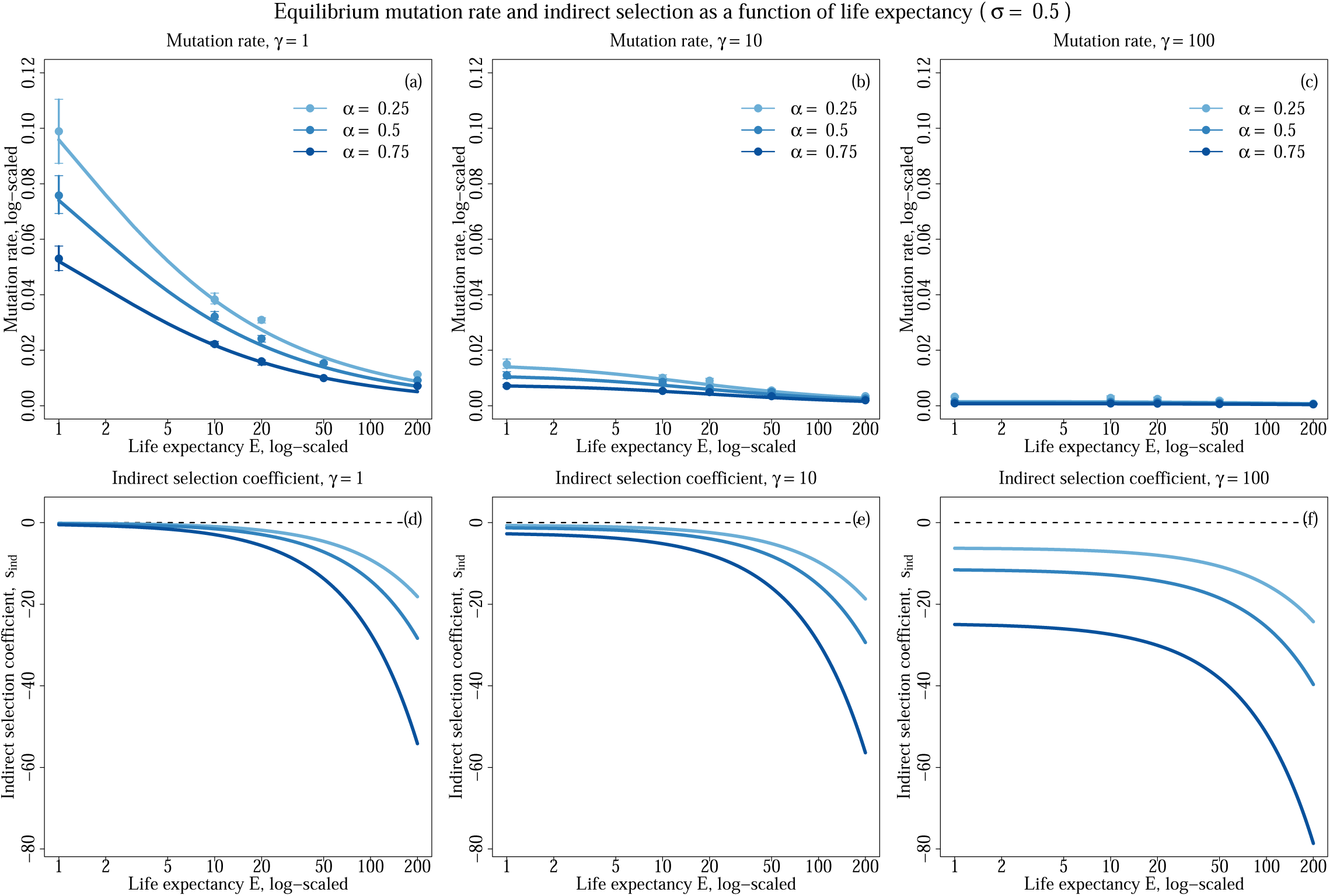
Evolutionarily stable mutation rate (top) and intensity of indirect selection (bottom) as a function of life expectancy (log-scaled) for various selfing rates (colors) and for *γ* = 1 (left), *γ* = 10 (middle) and *γ* = 100 (right). Other parameters values are *s* = 0.05, *h* = 0.3, *c* = 0.0014, *λ* = 20, and *σ* = 0.5. Dots depict simulation results and error bars depict the 95% confidence intervals. Lines depict analytical predictions.

The evolutionarily stable mutation rate decreases with life expectancy for all *γ* val-ues (Fig. 2a-c). In both cases, this is due to the greater number of opportunities to accumulate deleterious mutations in more long-lived species because they go trough more growth events, which in turn causes indirect selection to increase against alleles increasing the mutation rate because deleterious mutations become more numerous (Fig. 2d-f). Furthermore, the evolutionarily stable mutation rate is much lower when *γ* is larger, because increasing *γ* decreases the cost of replication fidelity (Equation 1), and increases the intensity of indirect selection on alleles increasing the mutation rate.

The mutation rate also decreases as the selfing rate (*α*) increases, which may seem counter-intuitive since selfing tends to reduce the number of deleterious mutations seg-regating in the population through purging Roze (2015). However, self-fertilisation also causes genetic associations between selected loci and the modifier to increase, thereby increasing indirect selection and resulting in a decrease of the evolutionarily stable mutation rate when the selfing rate increases as shown by Gervais and Roze (2017). The results presented in Fig. 2 were obtained assuming half of selfing events occured imperatively within the same section (*σ* = 0.5). Cases with *σ* = 0 and *σ* = 1 were also investigated and yielded very similar results, which are presented in Fig. S1 and S2, respectively, in Appendix II. We argue that the very small effect of *σ* on our results is due to the low evolutionarily stable mutation rate, which causes few somatic mutations to occur during growth, and to the fact that we assumed weak selection so that mutations have little effect on their bearer’s fitness.

### 3.2 Mutation-selection balance

Once the mutation rate has reached an equilibrium and the population has reached mutation-selection balance, we show in Appendix I.2.1 that a leading order approximation of the average number of mutations per haploid genome in juveniles (*n*) is given by

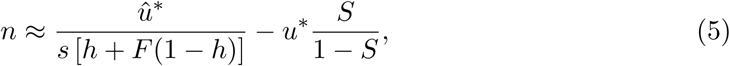

where 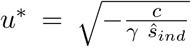, and 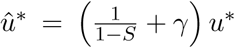 depicts the total mutation rate of the population over the course of one timestep, including both meiotic and somatic mutations. As for inbreeding depression, calculated between outcrossed and selfed individuals (*δ*_01_), it is given by

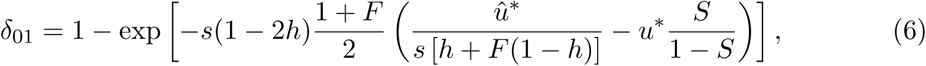

where 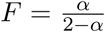, to leading order in *s*. Again, we neglect the impact of the proportion of selfing occuring within or between sections (*σ*) in our analytical work since it is negligible. Figure 3 shows the number of mutations per haploid genome among juveniles, (*n*, top row) and inbreeding and autogamy depression (*δ*_01_ and *δ*_12_, bottom row) at mutation-selection balance. Deviations between our analytical predictions (lines) and simulations results (dots) are observed. They can be explained by the slight differences between the predicted evolutionarily stable mutation rate and the equilibrium mutation rate reached by simulations, which build up large differences in *n* when life expectancy becomes high. Indeed, when the equilibrium mutation rate from the simulations is used to predict *n* instead of Equation (4), the agreement between predictions (open circles) and simulation results (dots) is restored.

**Figure 3:**
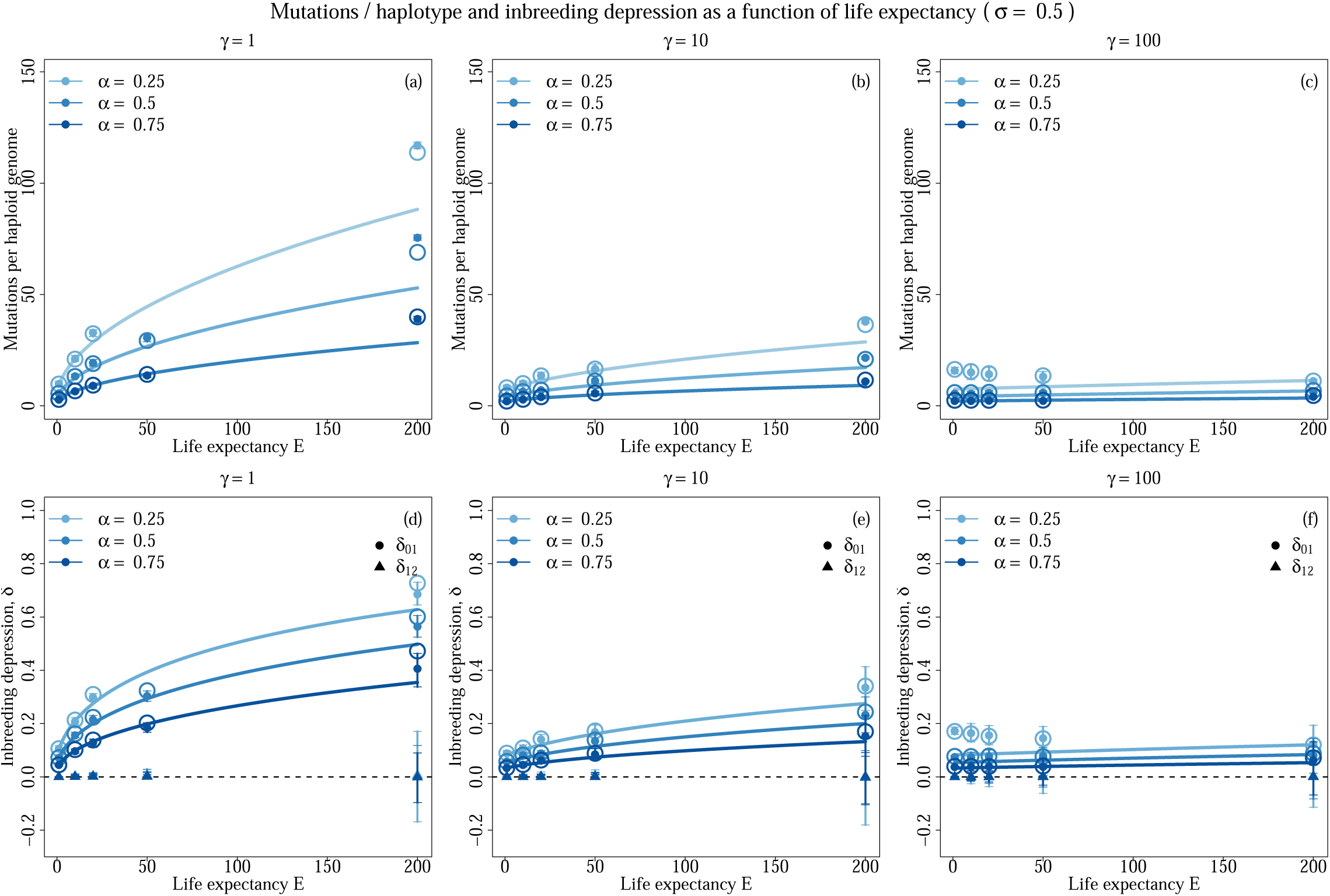
Average number of mutations per haploid genome (top) and inbreeding depression (bottom) as a function of life expectancy (log-scaled) for various selfing rates (colors) and *γ* = 1 (left), *γ* = 10 (middle) and *γ* = 100 (right). Other parameters values are *s* = 0.05, *h* = 0.3, *c* = 0.0014, *λ* = 20, and *σ* = 0.5. Filled dots depict simulation results and error bars depict the 95% confidence intervals. Lines depict analytical predictions. Open circles depict the value predicted by our analytical model when the equilibrium mutation rate from simulations is used instead of Equation 4. On the bottom row, dots indicate inbreeding depression (*δ*_01_), while triangles indicate autogamy depression (*δ*_12_).

The number of mutations maintained *n* increases as life expectancy increases in every case, due to the greater amount of opportunities for mutations to accumulate in more long-lived species. Indeed, in Equation (5), the denominator of the first term shows that the intensity of selection is independent of life expectancy, while the total mutation rate *û*^*∗*^ is a function of life expectancy. The increase of *n* with life expectancy becomes much lower when *γ* increases, to the point where it gets barely noticeable with *γ* = 100. Furthermore, *n* is lower when *γ* is higher despite the fact that many more mutations are produced during meiosis, because the evolutionarily stable mutation rate is much lower, so that the total mutation rate *û*^*∗*^ is lower (Fig. 2, Equation 5). As a result, inbreeding depression is lower when *γ* is higher, and increases when life expectancy increases, but this increase becomes less and less sharp as *γ* increases. Besides, *n* is lower when the selfing rate increases, as expected under weak selection (Roze, 2015). Furthermore, consistent with the negligible effect *σ* had on the evolution of the mutation rate, almost no autogamy depression is generated (triangles in Fig. 2, bottom row).

## 4 Discussion

### Evolution of the mutation rate

In this paper, we studied the evolution of the mutation rate when somatic mutations are assumed to be inheritable, as it is thought to be the case in plants (Scofield and Schultz, 2006; Lanfear, 2018). We showed that the evolutionarily stable mutation rate decreases as life expectancy increases because of the greater number of opportunities to accumulate mutations during growth in more long-lived species, which makes indirect selection against alleles increasing the mutation rate stronger. However, although the mutation rate per mutagenic event (*u*), that is per growth season or per meiosis in our model, decreased in more long-lived species, the total mutation rate (*û*), that is the rate at which mutations entered the population through both somatic growth and meiosis, increased. Hence, our results indicate that while we should expect more efficient mechanisms reducing the accumulation of deleterious mutations during growth to evolve in more long-lived species, so that their per unit of growth and per year mutation rate should be lower, their per generation mutation rates should still be higher. These predictions are in line with empirical evidence, which suggest that mutation rates per generation tend to be higher in more long-lived species although the mutation rates per unit of growth tend to be lower (Hofmeister et al., 2019; Hanlon et al., 2019; Orr et al., 2020).

We modeled the evolution of the mutation rate following the work of Kimura (1967), by assuming there is a direct fitness cost to DNA replication fidelity opposing the indirect selection generated by deleterious mutations linked to the modifier, so that the mutation rate was maintained greater than zero in response to a trade-off. An alternative mechanism, which is not mutually exclusive with the trade-off described above, was put forward by Lynch (2011). They proposed that selection should always act to reduce the mutation rate, down until it becomes so low that the selective advantage brought by any further reduction should be overwhelmed by genetic drift, thus maintaining non-zero mutation rates because alleles further decreasing the mutation rate should at some point become effectively neutral, thereby creating a lower bound for the evolution of the mutation rate (Lynch, 2011). This lower bound is inevitably influenced by effective population size, as it plays on the relative strength of selection and genetic drift. In our model, we overlooked Lynch (2011)’s lower bound by assuming a large and fixed population size. Yet, effective population sizes are expected to be higher in more long-lived species in which generations overlap (Felsenstein, 1971; Charlesworth, 1980; Petit and Hampe, 2006), which implies the lower bound described by Lynch (2011) should be met for lower mutation rates in said species. Hence, we expect the decrease in the evolutionarily stable mutation rate described in this study to become sharper in conditions where Lynch (2011)’s lower bound is expected to matter for the evolution of the mutation rate.

### Inbreeding depression

The larger total mutation rate in more long-lived species led to the maintenance of more mutations in the population at mutation-selection balance, and therefore to higher inbreeding depression in these species, consistent with results from meta-analyses which found inbreeding depression to increase in larger-statured, more long-lived species (Duminil et al., 2009; Angeloni et al., 2011). Importantly however, the magnitude of the increase in the total mutation rate, and therefore in inbreeding depression with life expectancy depended strongly on the relative mutagenicity of meiosis and growth, which was controlled by the *γ* parameter in our model. Indeed, while the increase in inbreeding depression was strong when *γ* was close to 1, that is when the same amount of mutation was produced during meiosis and during growth between two flowering seasons, it became smaller as *γ* increased, to the point of being barely noticeable for *γ* = 100. This was due to the decrease of the evolutionarily stable mutation rate as *γ* increased, which made the contribution of somatic mutations to the mutation load more and more negligible compared with meiotic mutations. Hence, according to our results, for somatic mutations to be the main driver of the empirically observed increase in inbreeding depression in more long-lived species, roughly the same amount of mutations should be produced during growth between two flowering seasons and during reproduction.

### Mating system evolution

Inbreeding depression is thought to be one of the main factors preventing the evolution of self-fertilisation (Lande and Schemske, 1985; Barrett and Harder, 2017). In Angiosperms, consistent with the observed increase in inbreeding depression in more long-lived species, there exists a strong correlation between mating systems and life-histories. Indeed, many self-fertilising species are annuals whereas most long-lived species are strictly outcrossing (Barrett and Harder, 1996; Munoz et al., 2016). Thus, somatic mutations accumulation was proposed as an explanation for this correlation (Scofield and Schultz, 2006). While our results indicate that inbreeding depression increases with respect to life expectancy due to somatic mutations accumulation, particularly when *γ* is small, this increase is tempered by the decrease of the evolutionarily stable mutation rate with life expectancy. Furthermore, in agreement with results obtained by Gervais and Roze (2017), we showed that the evolutionarily stable mutation rate decreases as the selfing rate increases because the modifier becomes more strongly associated with selected loci. These decreases of the mutation rate with respect to mating system and life expectancy, together with the purging effect of self-fertilisation (Roze, 2015), result in a substantial drop in the magnitude of inbreeding depression as the selfing rate increases in more long-lived species, potentially opening the way for the evolution of self-fertilisation. Hence, whether somatic mutations accumulation is sufficient to explain the correlation between life-history and mating system in Angiosperms when the mutation rate is allowed to evolve jointly with the mating system is an open question.

### Autogamy depression

In order to empirically estimate the contribution of somatic mutations accumulation to inbreeding depression using phenotypic data, a method was developed by Schultz and Scofield (2009). This method, called the autogamy depression test, relies on the comparison of the fitnesses of individuals produced by selfing within an inflorescence with those of individuals produced by selfing between distant inflorescences on the plant’s crown (Schultz and Scofield, 2009; Bobiwash et al., 2013). In our model, we performed such test by measuring autogamy depression (*δ*_12_). Contrary to inbreeding depression, we found autogamy depression to be almost null in every case, even in situations where the contribution of somatic mutations accumulation to inbreeding depression was high. This result can be explained by the low evolutionarily stable mutation rates, and by the fact that we only considered mutations with a weak fitness effect. It suggests that the autogamy depression test should only be able to detect mutations with a large fitness effect in large enough individuals, where mutations have had time to accumulate. Thus, it implies that detecting no autogamy depression in a given population cannot be taken as evidence of a negligible contribution of somatic mutations accumulation to the population’s mutation load.

### Mutagenicity of growth and meiosis

The results presented above suggest that valuable insights into the evolutionary relevance of somatic mutations and the evolution of the mutation rate in plants could be gained by further investigating the *γ* parameter in our model, which depicts the relative mutagenicity of meiosis and growth between two flowering seasons, and is likely influenced by at least three important factors that were unaccounted for in this study. First, it is necessarily influenced by how mutagenic meiotic divisions are in comparison with mitotic divisions, about which little is known although one may expect meiotic divisions to generate more mutations, as they generate many more double strand DNA breaks which are required for recombination and are known to be particularly mutagenic events (Magni and Von Borstel, 1962; Arbel-Eden and Simchen,2019). Second, it is influenced by the number of mitoses occurring between flowering seasons. This number depends on the growth habit of the considered species, because fast growing species undergo more mitoses per unit of time than slow growing species, and because the rate at which mitoses occur, and thus the growth rate, may interact with the evolution of the mutation rate. Indeed, investing in a higher fidelity of DNA replication may tend to slow down individual growth. Third, apart from mechanisms reducing the amount of mutations produced during growth, deleterious mutations may also be affected by intra-organismal selection, which may not only reduce the growth rate by eliminating mutated cells, but also efficiently purge deleterious mutations from the organism, so that little to no somatic mutation may be present in the gamete. This could in turn affect the evolution of the mutation rate. Little is known, however, about the actual efficacy of intra-organismal selection in removing deleterious mutations since it was seldom investigated theoretical (Otto and Orive, 1995), and mostly empirically demonstrated to occur in the case of strongly beneficial mutations (e.g. Edwards et al., 1990; Simberloff and Leppanen, 2019). The various elements discussed above show that *γ* is an emerging property of the interaction between a variety of mechanisms, which advocates for the development of theoretical models treating it as such rather than as a fixed parameter, by incorporating growth, mutation and selection at the cellular level.

## Acknowledgements

We would like thank Sylvain Billiard, Vincent Castric, Ludovic Maisonneuve, Denis Roze and Roman Stetsenko for taking the time to discuss this work and comment on the manuscript. This work was funded by the European Research Council (NOVEL project, grant #648321). The author also thanks the Région Hauts-de-France, and the Ministère de l’Enseignement Supérieur et de la Recherche (CPER Climibio), and the European Fund for Regional Economic Development for their financial support.

## APPENDICES

### I Derivation of analytical results

In this appendix, we detail the derivation of analytical results presented in the main text. The first section is dedicated to obtaining an approximation for the evolutionarily stable mutation rate, while the second is dedicated to the derivation of various population genetics quantities at mutation-selection balance.

#### I.1 Mutation rate evolution

In this section, we present the details of the method used to obtain the evolutionarily stable mutation rate approximation presented in the main text.

##### I.1.1 Defining age-dependent variables

###### Modifier locus

The mutation rate of an individual, given its genotype at the modifier, is given by

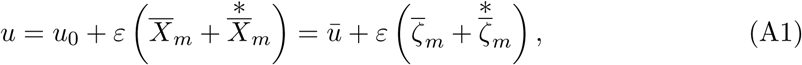

where *u*_0_ is the mutation rate coded by the resident allele, *ū* is the average mutation rate in the population, 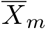 and 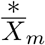 are indicator variables of the presence of the mutant allele on the paternally and maternally inherited chromosomes, respectively, and 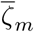 and 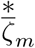 are their associated centered variables, with

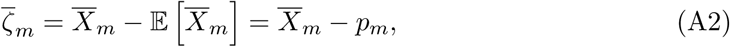

and respectively for 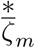 The bar in the notations indicates that these variables are considered over the whole population and not a particular age-class (see below). Importantly here, the genotype at the modifier of an individual does not vary between sections so that this notation is unnecessary in the above but is kept so that notations remain consistent all along derivations.

###### Selected loci

Since sections of various ages, that is sections bearing a different load of somatic mutations, contribute to reproduction, we define similar variables 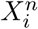 and 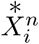 for the presence of deleterious mutations at the *i*^*th*^ selected locus, in the *n*^*th*^ section of individuals. Given the mutation rate *u* of an individual, its genotype in the (*n* + 1)^*th*^ section can be deduced from its genotype in the *n*^*th*^ section using Equation (A3):

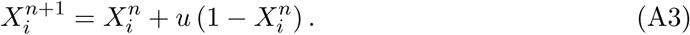

We can obtain the general term of this recursive sequence, which is given by

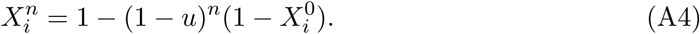

Injecting *ζ*−variables, developing *u* and rearranging Equation (A4), this yields,

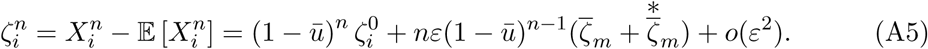

##### I.1.2 Summarising notations

For the sake of clarity, we summarise notations here. The same notations will be applied to indicator variables (*X*) and centered variables (*ζ*), allelic frequencies (*p*) and genetic associations (*D*, defined below).

###### Genetic associations

Let us forget for a moment the existence of age-classes in order to define properly genetic associations. Genetic associations are expectations of products of *ζ*-variables, and are in fact covariances between sets of indicator variables. In other words, they quantify the extent to which the frequency of sets of alleles at a given set of genetic positions deviate from the panmictic expectation, which assumes that alleles segregate independently both within and between loci. In general, we denote the genetic associations between the set of genetic positions 𝕌 on the paternal chromosome and the set of genetic positions 𝕍 on the maternal chromosome as

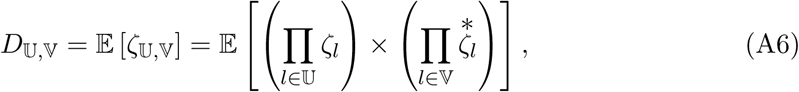

where the comma separates genetic positions situated on the paternal and maternal chromosomes. For instance, the association

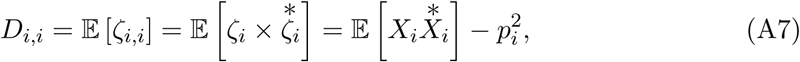

designates the excess in homozygotes at the *i*^*th*^ locus. Sets 𝕌 and 𝕍 can be empty, so that genetic associations between genetic positions all situated on the same chromosome (e.g. linkage disequilibrium) may be considered. For example, the association

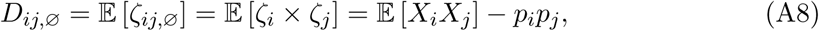

measures the linkage disequilibrium between loci *i* and *j* on the paternal chromosome. Since there is no separate sexes nor sex-specific effect in our model, we have

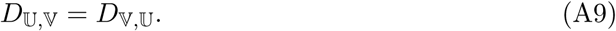

Hence, we define the condensed notation

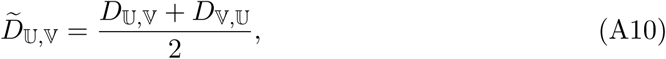

in order to shorten recursions.

###### Keeping track of age-classes and steps in the sequence of events

We stated above that 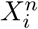(resp. 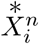) designates the indicator variable associated with the paternal (resp. maternal) position at the *i*^*th*^ selected locus in sections aged *n*. The same notation will be used for *ζ*-variables, genetic associations 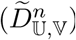 and allelic frequencies (e.g. 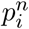). Importantly, while the genotype of sections at the modifier does not vary with age, selection may change allelic frequencies at the modifier differently in sections with different ages. Hence, the notation 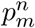 will also sometimes be used to indicate the fact that we consider allelic frequencies at the modifier among sections aged *n*. Hence, denoting *V* a generic variable (which may be *X, D, ζ* or *p*), and 𝕌 a generic set of genetic positions, 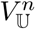indicates that the variable *V* is considered over the set 𝕌 of genetic positions among sections aged *n*. To keep track of the step in the sequence of events occurring over the course of one timestep we are looking at, that is for instance variables measured among parents, gametes or juveniles, we add an additional superscript separated from the age-class indication by a pipe. Denoting *k* a generic stage in the sequence of events, we thus have the notation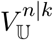. Finally, to indicate that we consider the average over all age-classes in the step *k* of the variable 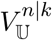, we will use the notation 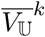.

##### I.1.3 Deriving an approximation for fecundity

We now derive an approximation for the fecundity of a section of age *n* as a function of its genotype, and relative to the entire parental population, that is sections of all ages. Let *W*_*n*_ be the fecundity of a section aged *n* and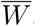, the fecundity of sections averaged over all ages. To leading order in ln*W*_*n*_ − ln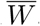, we have

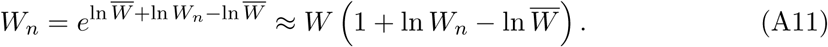

Furthermore, we have

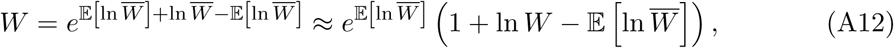

where 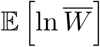 is the mean of the log-fitness. Thus, neglecting products of fitnesses, we have

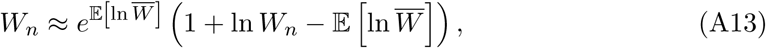

and since *W* is the average of *W*_*n*_ over all ages, the relative fecundity of a section aged *n* is given by

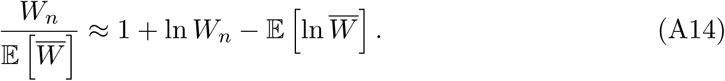

Let us now derive expressions for ln*W*_*n*_ and 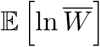. Expressed in terms of indicator variables, we have

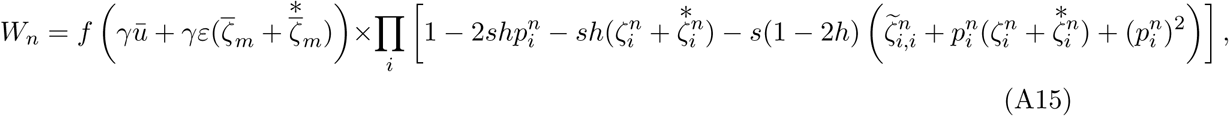

where *f* (*γu*) is the replication fidelity cost function, and *γ* allows one to tune the number of cell divisions occuring in meiosis relative to growth. Assuming selection to be weak and deleterious mutations to remain rare, so that terms in 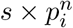 can be neglected, and using the fact that ln(1 + *x*) ≈ *x* when *x* is small, the log-fitness of a section of age *n* can be approximated as follows

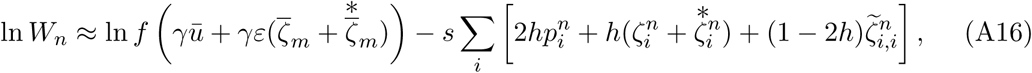

which to leading order in *ε* and in *s*, yields

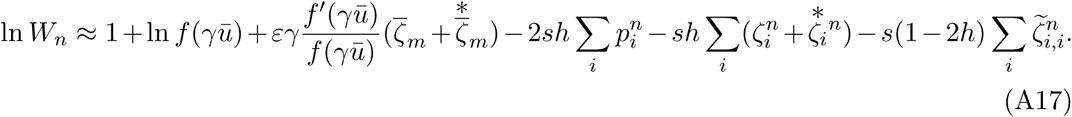

Since we have 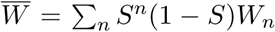, where *S* is the survival probability of individuals between mating events, we have

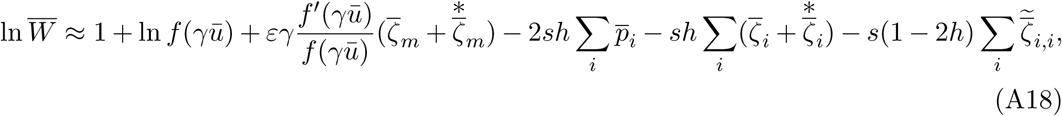

where the bar denotes the average over all ages, so that

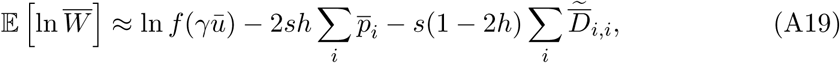

and,

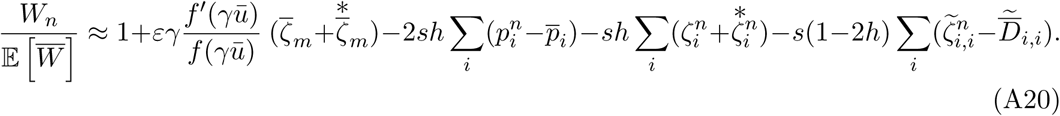

##### I.1.4 Computing the change in frequency at the modifier

Since we consider the fate of a rare mutant allele, the only relevant change in its frequency is due to selection. Assuming deleterious mutations remain at low frequencies, the frequency of the mutant in gametes produced by sections aged *n* following selection is given by

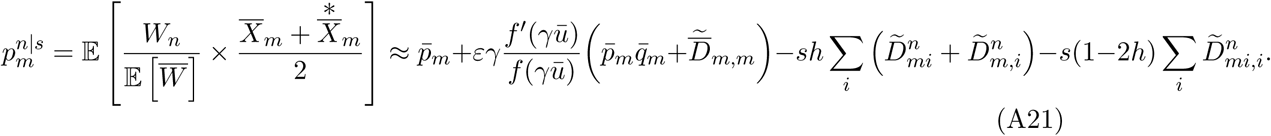

Thus, the frequency of the mutant among juveniles, which is the same as among gametes, is given by

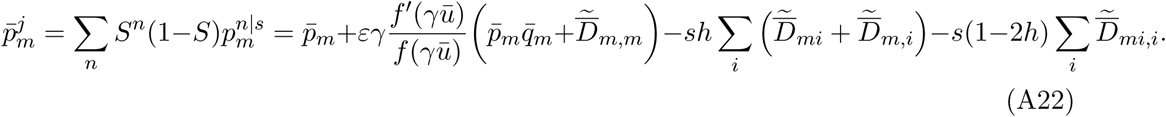

Since no selection occurs in parents, we have 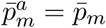, and

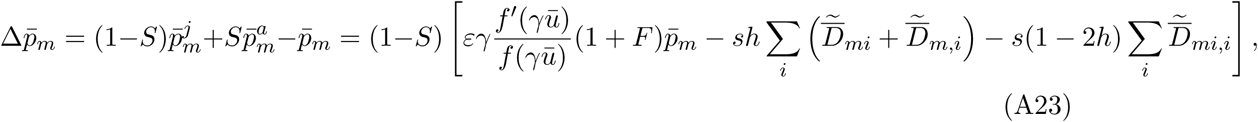

assuming *D*_*m,m*_ ≈ *Fp*_*m*_ (Roze, 2015). Equation (A23) can be broken down into two terms, one involving the cost function *f*, which depicts the direct selection acting on the modifier due to the replication fidelity cost, and a second term proportional to *s*, which depict the intensity of indirect selection acting on the modifier due to linked deleterious alleles.

##### I.1.5 Computing the indirect selection term

In order to obtain an approximation for the indirect selection term, which we denote *s*_*ind*_ following Gervais and Roze (2017), we need to obtain expressions for the genetic associations appearing in it at quasi-linkage equilibrium. On top of selection, these associations will also be affected by mutation, recombination and the mating system.

###### Two-way genetic associations

Let us begin with deriving the changes occuring in two-way genetic associations 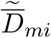 and 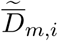 over one timestep. The effects of selection on such associations in gametes produced by sections aged *n* can be computed using

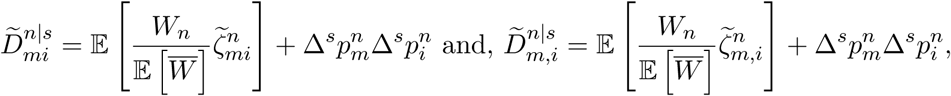

where Δ^*s*^*p*_*m*_ and Δ^*s*^*p*_*i*_ are changes in frequencies at the modifier and at the *i*^*th*^ selected locus, respectively. However, the product of changes in allelic frequencies is at best of order *ε* × *s*, so that we may neglect it in this context. Thus, we have

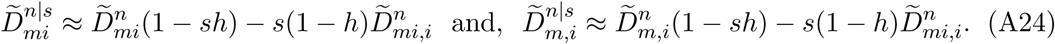

Selection is followed by meitoic mutation, which occurs at rate *γu*. Thus, in gametes we have

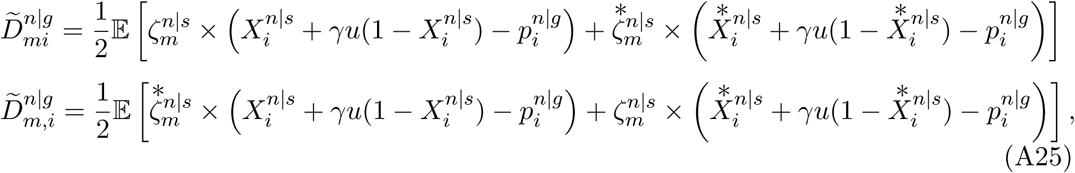

which to leading order in *ε* Yields

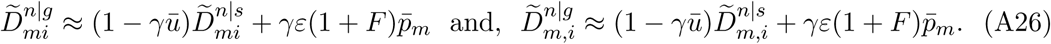

Reproduction occurs in three ways, outcrossing at rate 1 − *α*, selfing within section at rate *α*(1 − *σ*), and selfing between sections at rate *ασ*. While outcrossing does not generate genetic associations, both kinds of selfing do when associations involve genomic positions situated on different chromosomes. For the sake of simplicity, and because sections only differ from one another due to somatic mutations accumulation, which should be negligible at a given locus when the mutation rate is small, we will consider that selfing between and within sections have the same effect on genetic associations, so that in seeds following reproduction we have

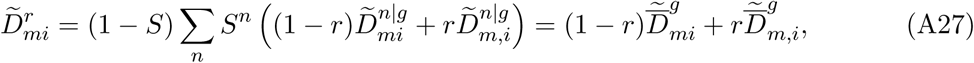

and,

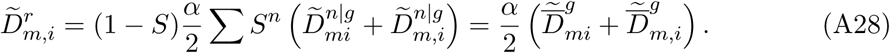

All individuals then undergo somatic mutation at rate *u*, which is incorporated in the exact same way as meiotic mutation was. Thus, we have

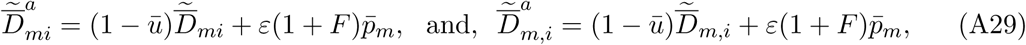

in adults and,

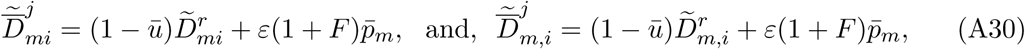

in juveniles.

###### Three-way association

Let us now turn to the derivation of the changes occuring the three-way association 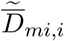 over the course of one timestep. Similar to other associations, we have,

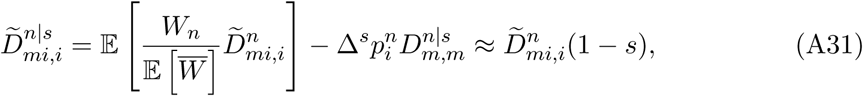

then, following the same method as in Equation (A25), we obtain

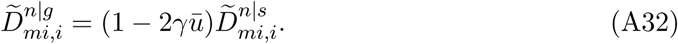

Among juveniles, we thus have

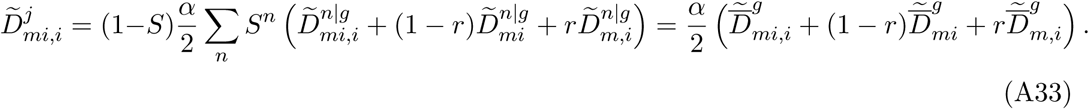

Among adults in the next timestep, we have

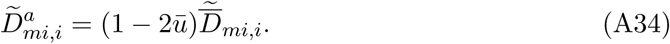

###### QLE expressions

To obtain expressions for genetic associations at quasi-linkage equilibrium, we need to solve Equation (A35) for 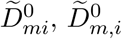, and 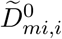

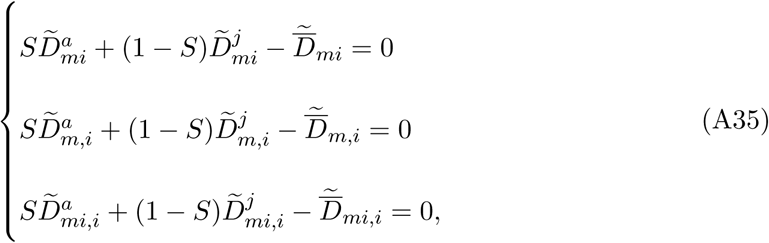

which, neglecting terms in ū and to leading order in *ε* and *s*, yields

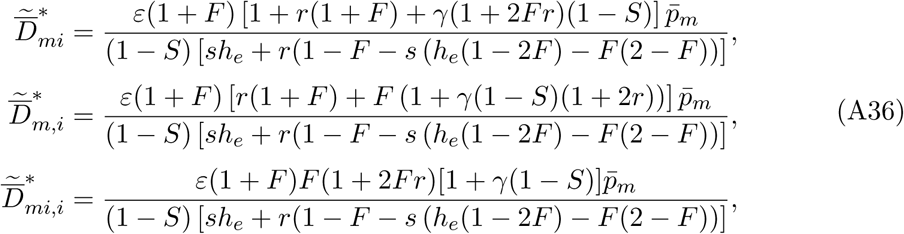

with *h*_*e*_ = *h* + *F* (1 − *h*).

###### Indirect selection term

Equation (A23) can be rearranged into

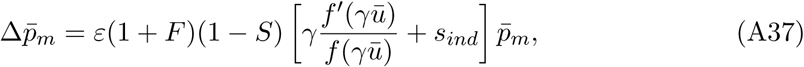

where the indirect selection term *s*_*ind*_ is given by

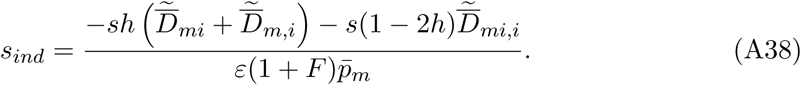

Using expressions presented in Equation (A36), this is

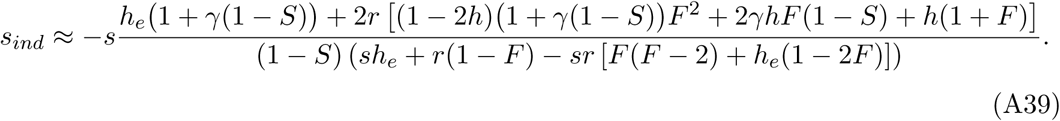

###### Integration over the genetic map

To account for the effect of the infinitely many loci present in the genome, one may integrate *s*_*ind*_ over the genetic map. The total length of the genetic map is *λ* (in cM), but since chromosomes are symetrical with respect to the centromere, we may focus one half of the genetic map, that is on the fraction of the chromosome situated between the centromere (*x* = 0), where the modifier is, and the tail of the chromosome 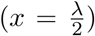. Thus, the probability that a crossing-over occurs between position 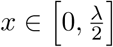 and the centromere, where the modifier is situated, is simply given by

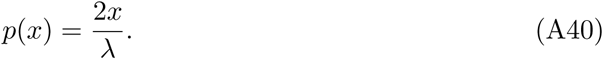

The number of crossing-overs during one meiosis event is drawn from a Poisson law with mean *λ*. Thus, the number of crossing-overs occuring on the 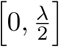 segment, *n*, follows a Poisson law with mean 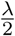 and we have

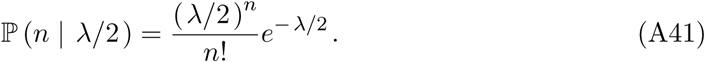

Then, the probability *k* of the *n* crossing-overs occur between position *x* and the centromere follows a Binomial law with probability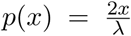. Importantly, recombination only effectively occurs when an uneven number of crossing-overs occurs on 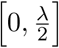, which given *n* occurs with probability

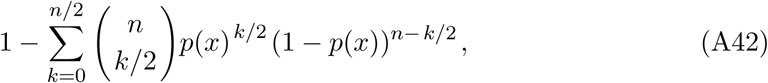

so that the probability of recombination effectively occurring between position x and the centromere along 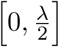 is given by

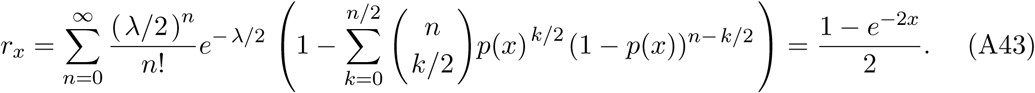

Plugging Equation (A43) in Equation (A39), that is swapping *r* for *r*_*x*_, the indirect selection term accounting for the genetic map is given by

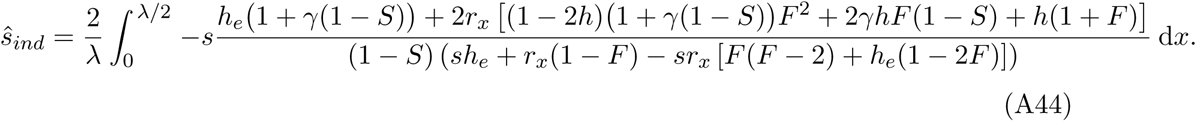

Thus, using *f* (*γu*) cost function described in the main text, the change in frequency at the modifier can be written as

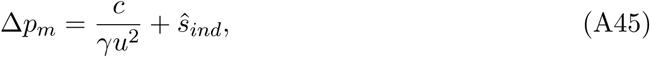

so that the evolutionarily stable mutation rate is given by

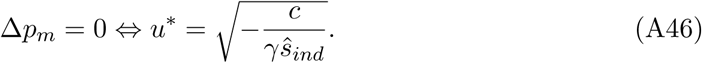

### I.2 Mutation-selection balance

Once the population has reached its equilibrium mutation rate, it reaches mutation-selection equilibrium. The average number of mutations per haploid genome and the resulting inbreeding depression can then be obtained by assuming the mutation rate is constant.

#### I.2.1 Average number of mutations per haploid genome

##### Among juveniles

Using (Equation A20) in Appendix I.2, and assuming the modifier is fixed at the evolutionarily stable mutation rate (*i*.*e. u* = *u*^*∗*^ and *ε* = 0), the frequency of the deleterious allele at the *i*^*th*^ selected locus among gametes produced sections aged *n* due to selection is given by

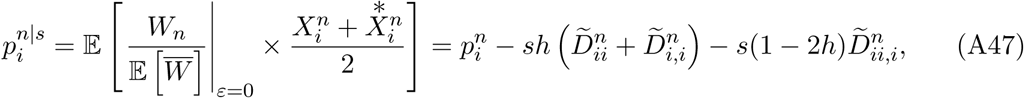

which noting that

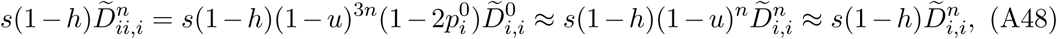

and that,

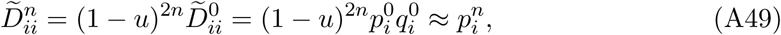

can be rearranged into

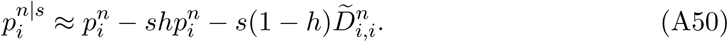

Thus, in juveniles following meiotic and somatic mutation with have

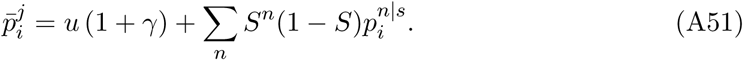

As for the excess in homozygotes among juveniles, we neglect the effects of selection and mutation so that we have

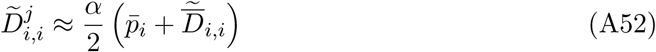

##### Among adults

Among adults, selection does not have any effect, so that we simply have

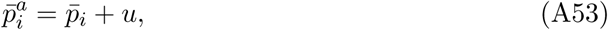

and,

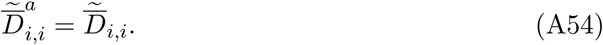

##### Equilibrium excess in homozygotes

Neglecting the effects of mutation and selection on homozygosity at selected loci, we have

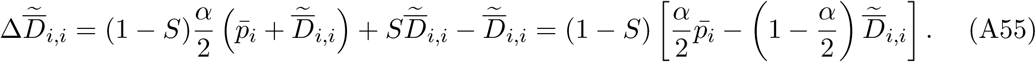

Thus, the equilibrium excess in homozygotes at selected loci is given by

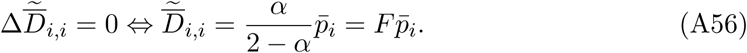

##### Equilibrium number of mutations

The change in frequency of the deleterious allele at the *i*^*th*^ locus is given by

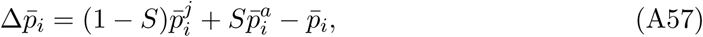

so that at equilibrium, with 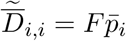, we have

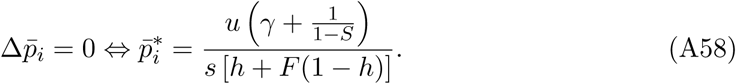

In simulations, we measure the average number of mutations per haploid genome in seeds, which can be obtained as follows:

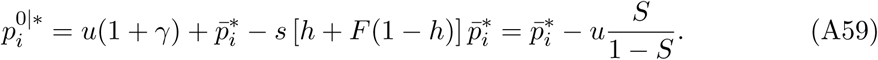

#### I.2.2 Inbreeding depression

In this paper, we chose to measure inbreeding depression at the scale of whole individuals even though individuals are chimeric from the genetic point of view, in order to remain consistent with its classical definition. To obtain an analytical approximation for inbreeding depression, we thus need to compute the mean lifetime reproductive success of selfed and outcrossed individuals.

##### Lifetimes fitness expression

Since mutations have a null probability of occurring on both alleles at the same time at a locus, so that homozygotes are never created by mutation, the fecundity of a section aged *i* can be approximated as

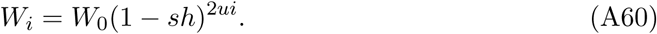

Thus, the total fecundity of an individual aged *k* is

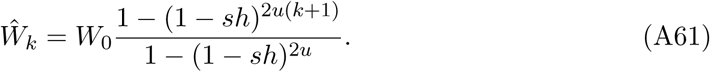

Hence, lifetime fitness can be computed as

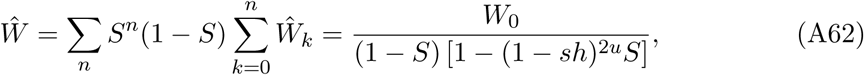

##### Inbreeding depression

Using Equation (A62), inbreeding depression can be expressed as

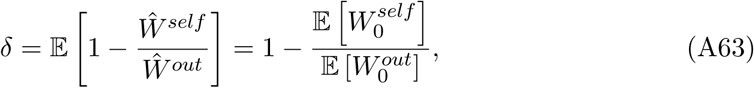

where *Ŵ*^*self*^ and *Ŵ*^*out*^ are lifetime fitness measured among the selfed and the outcrossed, respectively. We now need to obtain leading order approximations for 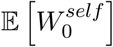 and 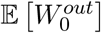. For any subpopulation *sub*, we have

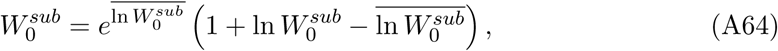

to leading order in ln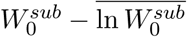, so that

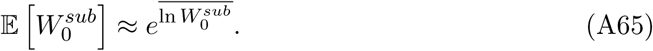

Furthermore, from Equation (A16), we have

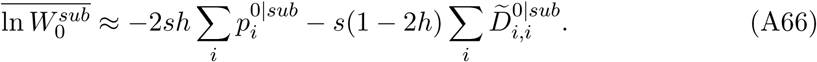

In the case of selfed and outcrossed individuals, we have 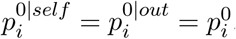. Besides, the excess in homozygotes among outcrossed individuals is 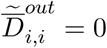, while among the selfed, it is given by 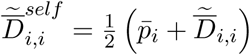 Thus, we have

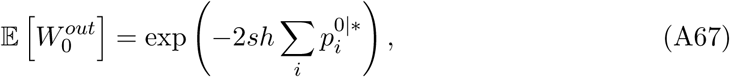

and,

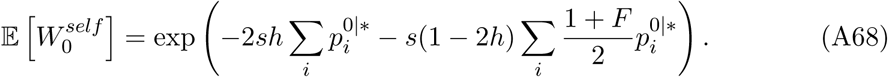

Hence, inbreeding depression in our model is given by

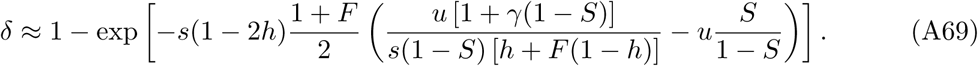

### II Additional graphs

**Figure S1:**
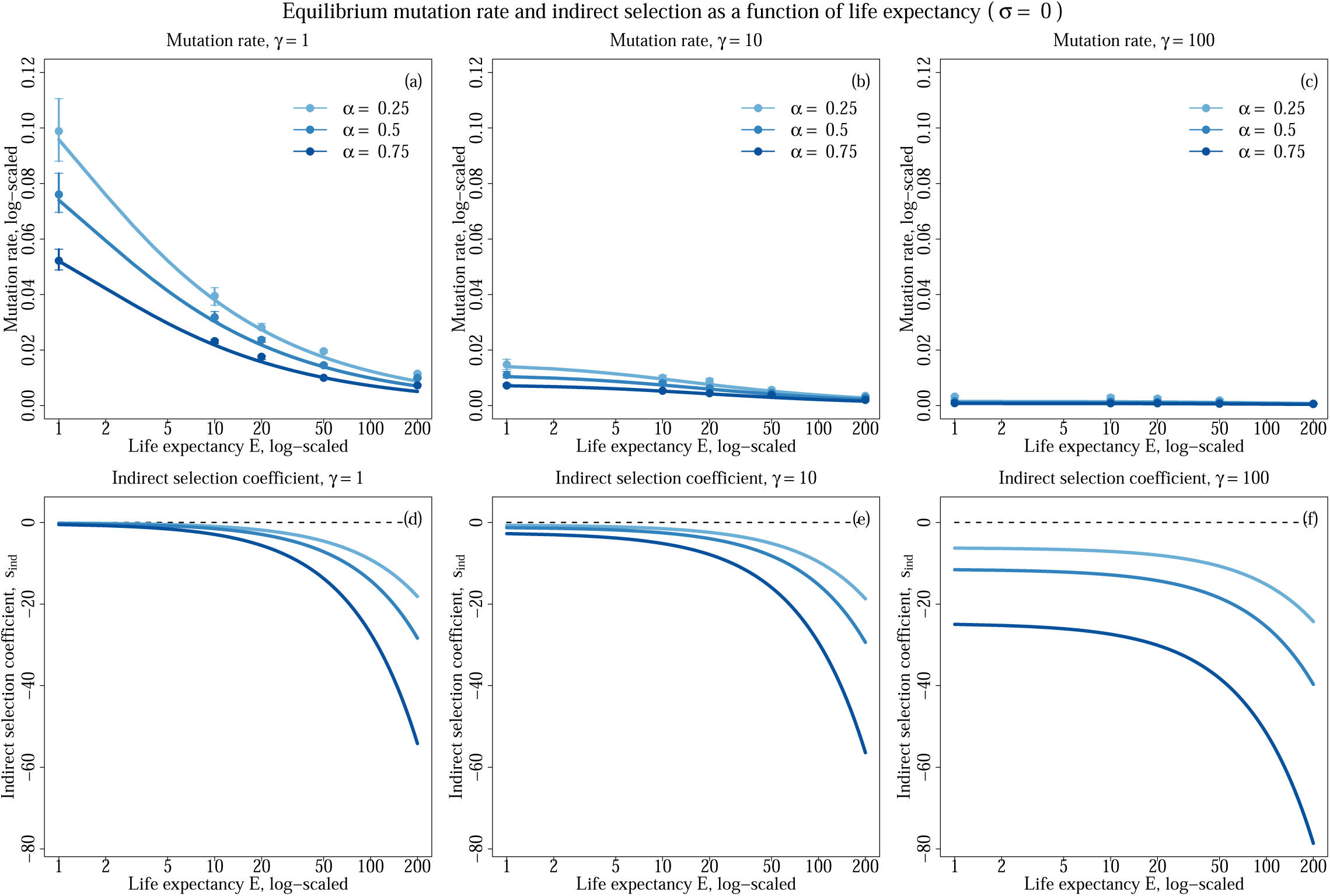
Evolutionarily stable mutation rate (top) and intensity of indirect selection (bottom) as a function of life expectancy (log-scaled) for various selfing rates (colors) and for *γ* = 1 (left) and *γ* = 10 (right). Other parameters values are *s* = 0.05, *h* = 0.3, *c* = 0.0014, *λ* = 20, and *σ* = 0. Dots depict simulation results and error bars depict the 95% confidence intervals. Lines depict analytical predictions.

**Figure S2:**
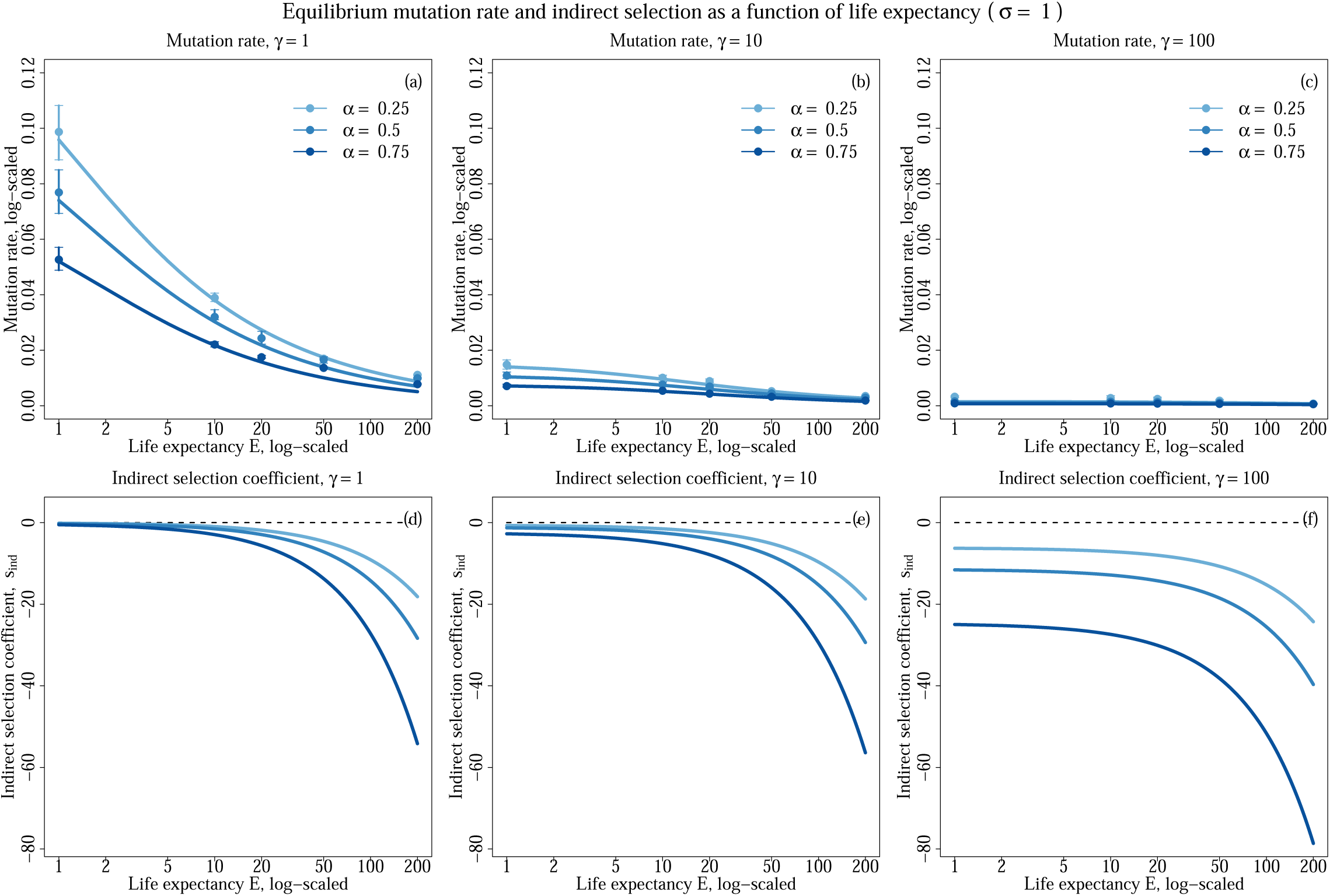
Evolutionarily stable mutation rate (top) and intensity of indirect selection (bottom) as a function of life expectancy (log-scaled) for various selfing rates (colors) and for *γ* = 1 (left) and *γ* = 10 (right). Other parameters values are *s* = 0.05, *h* = 0.3, *c* = 0.0014, *λ* = 20, and *σ* = 1. Dots depict simulation results and error bars depict the 95% confidence intervals. Lines depict analytical predictions.

**Figure S3:**
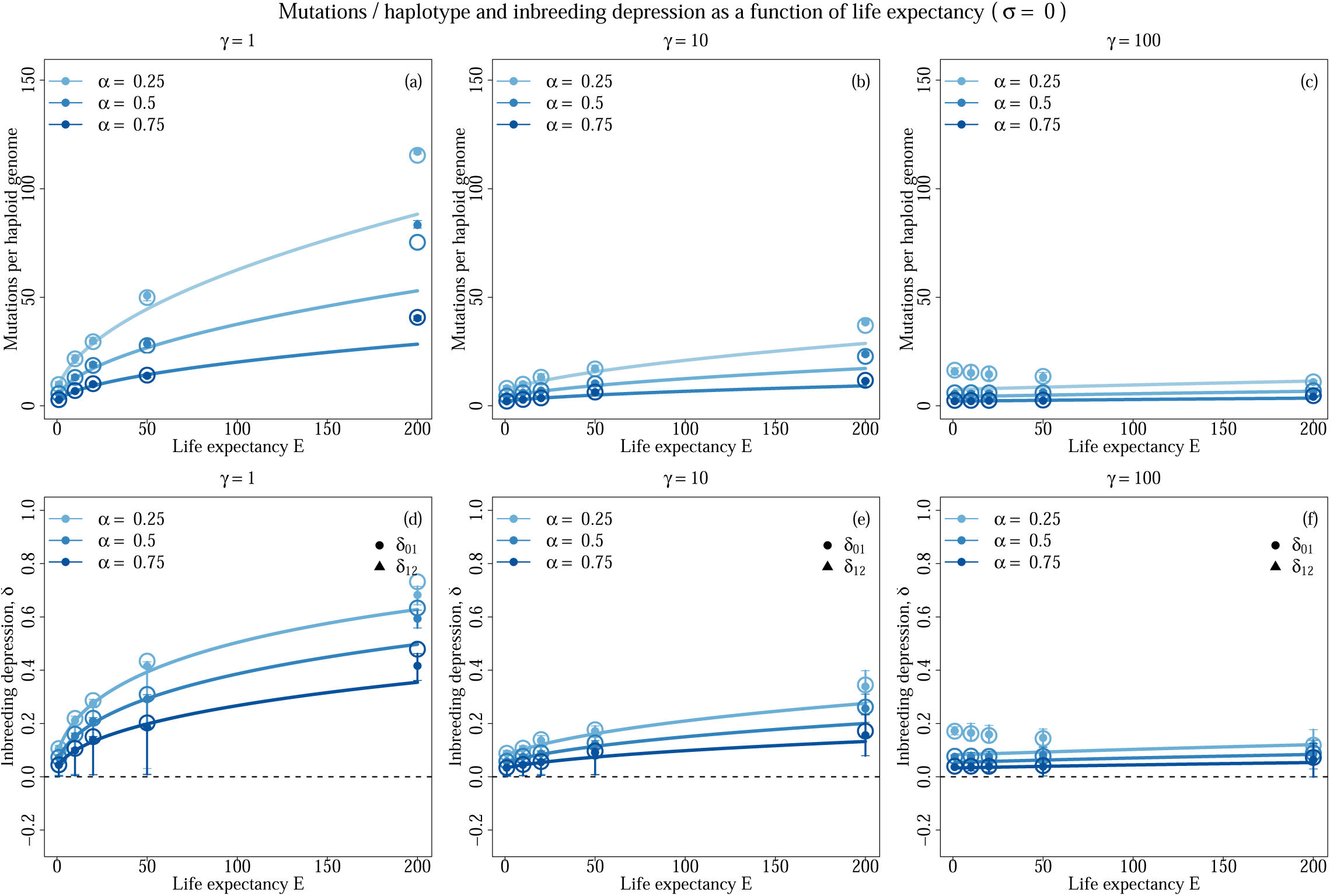
Average number of mutations per haploid genome (top) and inbreeding depression (bottom) as a function of life expectancy (log-scaled) for various selfing rates (colors) and for *γ* = 1 (left) and *γ* = 10 (right). Other parameters values are *s* = 0.05, *h* = 0.3, *c* = 0.0014, *λ* = 20, and *σ* = 0. Filled dots depict simulation results and error bars depict the 95% confidence intervals. Lines depict analytical predictions. Open circles depict the value predicted by our analytical model when the equilibrium mutation rate from simulations is used instead of Equation 4. On the bottom row, dots indicate inbreeding depression (*δ*_01_), while triangles indicate autogamy depression (*δ*_12_).

**Figure S4:**
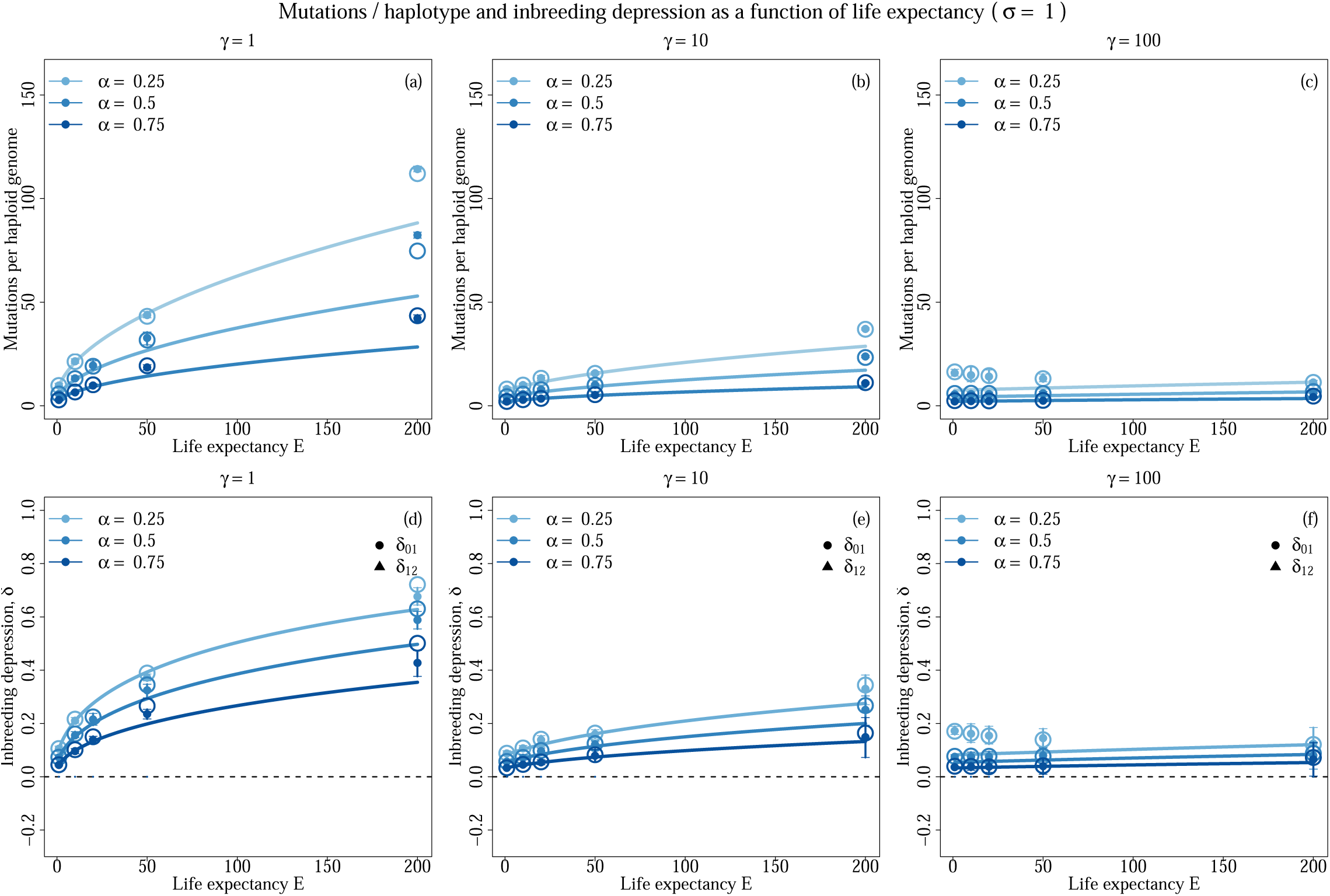
Average number of mutations per haploid genome (top) and inbreeding depression (bottom) as a function of life expectancy (log-scaled) for various selfing rates (colors) and for *γ* = 1 (left) and *γ* = 10 (right). Other parameters values are *s* = 0.05, *h* = 0.3, *c* = 0.0014, *λ* = 20, and *σ* = 1. Filled dots depict simulation results and error bars depict the 95% confidence intervals. Lines depict analytical predictions. Open circles depict the value predicted by our analytical model when the equilibrium mutation rate from simulations is used instead of Equation 4. On the bottom row, dots indicate inbreeding depression (*δ*_01_), while triangles indicate autogamy depression (*δ*_12_).

